# Seedling recruitment after fire: Disentangling the roles of microsite conditions and seed availability

**DOI:** 10.1101/2024.11.12.623237

**Authors:** Jared J. Beck, Stuart Wagenius

**Author notes:** Corresponding author –.

## Abstract

Periodic fire enhances seedling recruitment for many plant species in historically fire-dependent ecosystems. Fire is expected to promote recruitment by generating environmental conditions that promote seedling emergence and survival. However, fire may also increase flowering and seed production. This makes it difficult to distinguish the effects of microsite conditions from seed availability in observational studies of seedling recruitment. Experiments that manipulate seed inputs across a representative range of conditions are needed to elucidate how seed availability versus microsite conditions influence post-fire seedling recruitment and plant demography. We experimentally manipulated time since fire across 36 patches of remnant tallgrass prairie distributed across 6400 ha in western Minnesota (USA). Over two years, we sowed 11,057 *Echinacea angustifolia* (Asteraceae) seeds across 84 randomly placed transects and tracked 974 experimentally sown seedlings to evaluate how time since fire influenced seedling emergence and survival after experimentally controlling for variation in seed inputs. We also quantified six environmental variables and evaluated whether these covariates were associated with seedling emergence and survival. Fire influenced both seedling emergence and seedling survival. Seedlings emerged from approximately 1 percent of all seeds sown prior to experimental burns. Seeds sown one year after experimental burns emerged at 15 times the rate of seeds sown in the fall before burns, but emergence then declined as time since fire increased. Sowing seeds at high densities reduced rates of seedling emergence but increased overall recruitment. Increases in litter depth were associated with reduced emergence. Meanwhile, the probability that seedlings survived to late summer was greatest when they emerged 0-1 years after fire. The probability of seedling survival decreased with litter depth and increased with the local density of conspecific seedlings. Our findings experimentally support widespread predictions that fire enhances seedling recruitment by generating microsite conditions favorable for seedling emergence and survival – especially by increasing the light available to newly emerged seedlings. Nevertheless, recruitment also increased with seed inputs indicating that both seed availability and microsite conditions influence post-fire recruitment. Explicitly discriminating between seed-limitation and microsite-limitation is critical for understanding the demographic processes that influence plant population dynamics in historically fire-dependent ecosystems.

## INTRODUCTION

Fire influences many aspects of the life history and demography of plants across historically fire-dependent ecosystems worldwide (Bond & Keeley, 2005; L. T. Kelly et al., 2020; Pausas & Keeley, 2014). For many plant species in these systems, periodic fire is expected to promote seedling recruitment by generating conditions favorable to the growth and survival of seedlings (Lamont & Downes, 2011; Leach & Givnish, 1996a; Menges & Dolan, 1998; Menges & Kimmich, 1996; Pausas & Keeley, 2014; Satterthwaite et al., 2002; Tyler, 1995). Enhanced seedling recruitment after fire can promote the growth and persistence of plant populations (e.g., Leach & Givnish, 1996; Menges & Dolan, 1998; Nordstrom et al., 2021) and may reflect various traits and life history characteristics that have evolved in the context of frequent fire (e.g., Beck et al., 2024; Lamont & Downes, 2011; Pausas & Keeley, 2014). While the beneficial effects of fire on seedling recruitment are often attributed to post-fire environmental conditions that promote seedling growth and survival (i.e., favorable microsite), fire may influence three distinct processes that contribute to variation in seedling recruitment: seed production, seedling emergence, and seedling survival. Directly quantifying the extent to which enhanced seedling recruitment after fire reflects greater seed production and more seeds entering the seed bank compared to improved microsite conditions that foster seedling emergence and survival seedling emergence and survival (i.e., seed-limitation versus microsite-limitation sensu Crawley, 1990; Eriksson & Ehrlen, 1992; Turnbull et al., 2000) is essential for elucidating the mechanisms by which fire influences seedling recruitment. Studies capable of generalizing these mechanistic inferences about seed-versus microsite-limitation across a representative sample of populations can offer new insights about plant population and community responses to fire as well as the evolutionary ecology of plants within historically fire-dependent systems.

Observed rates of seedling recruitment in natural populations reflect the cumulative effects of variation in the number of seeds that enter the seed pool and survive through dormancy, the germination and growth of seeds (i.e., seedling emergence), and seedling survival after emergence (Crawley, 1990; Eriksson & Ehrlen, 1992; Turnbull et al., 2000). Fire has the potential to influence each of these distinct processes in historically fire-dependent ecosystems (e.g., Lamont & Downes, 2011; Tyler, 1995). For instance, fire exposes mineral soil, which improves seed-soil contact (e.g., McConnell & Menges, 2002) and increases the short-term availability of several important macronutrients (e.g., Ojima et al., 1994). Fire can also suppress woody plants and other dominant species that may competitively exclude seedlings (e.g., Peterson & Reich, 2007). Perhaps most importantly, fire generally increases the light available to newly emerged seedlings by consuming accumulated leaf litter and other plant material (e.g., Bond & Keeley, 2005). Increased light availability could enhance seedling growth and survival (Alstad et al., 2016; L. Zirondi et al., 2021; Leach & Givnish, 1996; McConnell & Menges, 2002). For these reasons, fire is widely expected to influence environmental conditions in ways that promote seedling emergence and seedling survival. In other words, fire is hypothesized to promote seedling recruitment by ameliorating microsite limitation. However, fire also stimulates flowering in many plant species across historically fire-dependent ecosystems (Beck et al., 2024; Fidelis & Zirondi, 2021; Lamont & Downes, 2011). This synchronized post-fire flowering is often associated with enhanced seed production (Beck et al., 2023; Richardson & Wagenius, 2022; Vickery, 2002; Wagenius et al., 2020). Thus, increased seedling recruitment after fire could reflect the greater seed inputs resulting from mass flowering, post-fire environmental conditions that favor greater seedling emergence, or factors that promote seedling survival after emergence.

Differentiating between the influences of seed availability compared to microsite conditions on post-fire seedling recruitment is critical for elucidating demographic processes that promote population growth and persistence in historically fire-dependent ecosystems such as North American tallgrass prairie. Fire played an important historical role in shaping the habitat structure and biological diversity of these once expansive grasslands (Anderson, 2006; Axelrod, 1985; Curtis, 1959; Gleason, 1913). Fires occurred at an estimated frequency of 1- to 5-year intervals reflecting a combination of lightning-caused conflagrations and the active use of fire by indigenous groups (Abrams, 1985; Allen & Palmer, 2011; Anderson, 1990; Stewart, 2009). The destruction and fragmentation of habitat coupled with intentional fire suppression over the past 100-200 years has substantially reduced the frequency of fire in most tallgrass prairie remnants (Curtis, 1959; Leach & Givnish, 1996; McClain et al., 2021; Umbanhowar, 1996). With fire now infrequent, native plant diversity within prairies has declined rapidly (Alstad et al., 2016; Bowles & Jones, 2013; Leach & Givnish, 1996). While beneficial effects of periodic fire on plant diversity in prairies are often attributed to enhanced seedling recruitment (Alstad et al., 2016; Leach & Givnish, 1996), two distinct demographic mechanisms could contribute to greater seedling recruitment and population persistence: improved seedling emergence and survival due to altered microsite conditions (Leach & Givnish, 1996) and greater seed inputs to the seed bank due to enhanced flowering and seed production (Wagenius et al., 2020).

Observational studies of seedling recruitment in prairies commonly report greater seedling recruitment after fire (Benson & Hartnett, 2006; Glenn-Lewin et al., 1990; Menges & Dolan, 1998; Nordstrom et al., 2021), but such studies cannot discriminate the roles of availability of seeds and of microsites (Crawley, 1990; Eriksson & Ehrlen, 1992). Seed addition experiments overcome many of the limitations inherent to observational studies of seedling recruitment. By manipulating the number of seeds entering the seed bank and quantifying establishment of experimentally sown seed, seed addition experiments can effectively distinguish effects of seed and microsite availability on recruitment (Crawley 1990; Eriksson & Ehrlen, 1992; Setterfield, 2002). These experiments provide evidence of seed-limitation when increases in seed availability result in greater seedling recruitment. Meanwhile, these experiments provide evidence of microsite-limitation when recruitment depends on environmental conditions after controlling for variation in seed inputs. Note that seed availability and microsite conditions can co-limit recruitment (e.g., Eriksson & Ehrlen, 1992). Furthermore, the resulting variation in conspecific seedling density offers the opportunity to gain additional insights into density-dependent seedling survival, to the extent the variation is independent of underlying environmental conditions.

Previous seed addition experiments in tallgrass prairie suggest that post-fire microsite characteristics exert a strong influence on seedling emergence and seedling survival independent of seed input (Alstad et al., 2018; Wagenius et al., 2012). However, it remains unclear whether findings from previous experimental studies can be generalized across the broad range of conditions encountered by seedlings in natural populations. Previous seed addition experiments related to fire have been conducted at few sites (e.g., Alstad et al., 2018), within homogeneous restorations, or, in locations not representative of conditions in natural populations (e.g., Wagenius et al., 2012). All of these issues complicate extrapolation to natural plant populations because natural populations span considerable variation in biotic and abiotic environmental conditions (Quintana-Ascencio, 2023). Furthermore, previous work in prairies and other fire-dependent habitats has shown that fire can exert context-dependent effects on seedling recruitment (Iacona et al., 2010; Menges & Hawkes, 1998; Myers & Harms, 2011; Wagenius et al., 2012). Spatial and temporal replication is needed to assess how the magnitude of fire effects compares to underlying variation in seedling vital rates in natural populations. The relative magnitude of sources of variation is critical for assessing fire effects on seedling recruitment and subsequent effects on plant demography (Nordstrom et al, 2021).

We therefore conducted a seed addition experiment across 36 remnant patches of tallgrass prairie to experimentally assess how time since fire influences seedling emergence and seedling survival in natural populations. Furthermore, we sought to evaluate how seedling emergence and survival were related to seed availability, conspecific seedling density, and several abiotic aspects of the environment. Our experiment focused on *Echinacea angustifolia* (Asteraceae) – a long-lived herbaceous species widespread across central North American grasslands. Previous studies investigating the demography of *E. angustifolia* seedlings have yielded insights into seedling dynamics and their implications for plant demography. Long-term observational studies of seedling recruitment in natural populations showed considerable variation within and among populations as well as among years (Richardson et al., 2024; Waananen et al., 2024). Using observational recruitment data encompassing a limited set of prescribed burns, Nordstrom et al. (2021) found seedling recruitment (combined effects of seed availability, seedling emergence, and seedling survival in the first year) tended to increase after fire which strongly influenced demographic rates. In a seed addition experiment, Wagenius et al. (2012) found evidence that fire benefits seedling emergence and survival in old fields and prairie restorations (i.e., not remnant prairie).

Our experiment here builds on this previous work in several important ways. First, we conducted our experiment within natural populations and use random placement of transects within sites to ensure valid inferences across natural populations. Second, experimental seed addition enables us to control and manipulate variation in the number of seeds entering the seed bank. This experimental manipulation of seed availability and our explicit focus on experimental seedlings (i.e., we do not assess natural recruitment) enable us to discriminate between seed-limitation and microsite-limitation. Third, we experimentally manipulated time-since-fire across 36 prairie remnants over five years resulting in true spatial and temporal replication of burn treatments across representative conditions. Finally, we quantified the relationship between emergence and survival with six microsite characteristics that may mediate or influence seedling responses to fire: local conspecific seedling density, light availability, litter depth, soil compaction, slope, and aspect. Building directly on these methodological advances, we address four questions in this study: (1) How does time since fire influence seedling emergence in natural populations? (2) To what extent does variation in seedling emergence reflect environmental conditions versus the density of experimentally sown seeds? (3) How does time since fire influence seedling survival? And (4) does variation in seedling survival reflect environmental factors?

## MATERIALS AND METHODS

### Study species

Our experimental investigation of fire effects on seedling recruitment focused on *Echinacea angustifolia* (Asteraceae). This herbaceous perennial is native to central North America and widespread across grasslands west of the Mississippi River. The species inhabits dry prairie hills and locations with well-drained soils. Individual plants produce a deep taproot and can live for decades. From seed, plants do not flower for the first time before 5 years of age and many individuals take more than a decade (Richardson et al., 2024). Plants do not spread vegetatively and reproduce only via seed. Sexually mature plants do not flower every year. In a year when an *E. angustifolia* individual flowers, it typically produces one composite head encompassing ca. 150 florets, though a flowering plant can produce as many as 29 heads. Each uniovulate floret yields a dry fruit (an achene) that may or may not contain an embryo (i.e. a seed). A floret must receive compatible pollen to produce an embryo. Achenes are produced regardless of success of pollination. Achenes, which ripen and fall to the ground in fall (September – November), have no specialized dispersal mechanism. Seeds germinate in spring after overwintering and do not persist in a seedbank. In greenhouse conditions, light enhances germination rates (Feghahati & Reese, 1994). Seedlings emerge late spring (May) to early summer (June) and produce a single true leaf, rarely two.

### Study area

Our experiment encompassed 36 patches of tallgrass prairie distributed across a 6400 ha study area in western Minnesota (USA) centered near 45°49’ N, 95°43 W (Fig. S1). All focal sites support extant populations of *E. angustifolia.* Long-term studies of plant demography have been conducted at most (28 of 36) sites continuously since 1996. The study area comprises a primarily agricultural landscape with scattered patches of remnant tallgrass prairie. Sites range from ca. 0.1 hectares (ha) to 10 ha. These focal sites span the breadth of habitat occupied by *E. angustfolia* – ranging from gravelly roadsides and railroad rights-of-way to dry prairie hills – and encompass variation from high-quality prairie preserves with high native plant diversity to degraded remnants with low native plant diversity.

### Experimental design

We established 84 transects across the 36 focal sites (Fig. 1). The location of each 4 m long transect was selected using a stratified random sample of *E. angustifolia* patches within focal sites. In most sites, transect locations were selected at random after defining the spatial extent occupied by *E. angustifolia.* In several sites that encompassed distinct clusters of *E. angustifolia*, we first defined the spatial extent of each cluster before randomly selecting transect locations within each cluster to obtain a stratified random sample. These procedures for selecting transect locations are critical for obtaining an unbiased sample of locations as a basis for generalizable inferences about seedling recruitment dynamics within populations. The number of transects assigned to each site was proportional to site area. Each 4 m long transect was partitioned into four 1 m segments with nails permanently marking the end points of each transect (Fig. S1). We manually removed any *E. angustifolia* seed heads near (i.e., within 0.5 m) experimental transects during the summer prior to sowing to avoid including natural seedling recruits in our surveys. In previous observational studies of seedling recruitment *in E. angustifolia*, natural recruits were very rarely found more than 0.5 m from the maternal plant (Richardson et al., 2024). Moreover, the density of newly emerged natural recruits from a random sample of locations in our study area was estimated to be 0.02 seedlings per m^2^. The manual removal of nearby heads, the limited dispersal of *E. angustifolia* seeds, and the low density of natural recruits in these remnants make us confident that natural recruitment had little if any influence on our experiment.

**Fig. 1.**
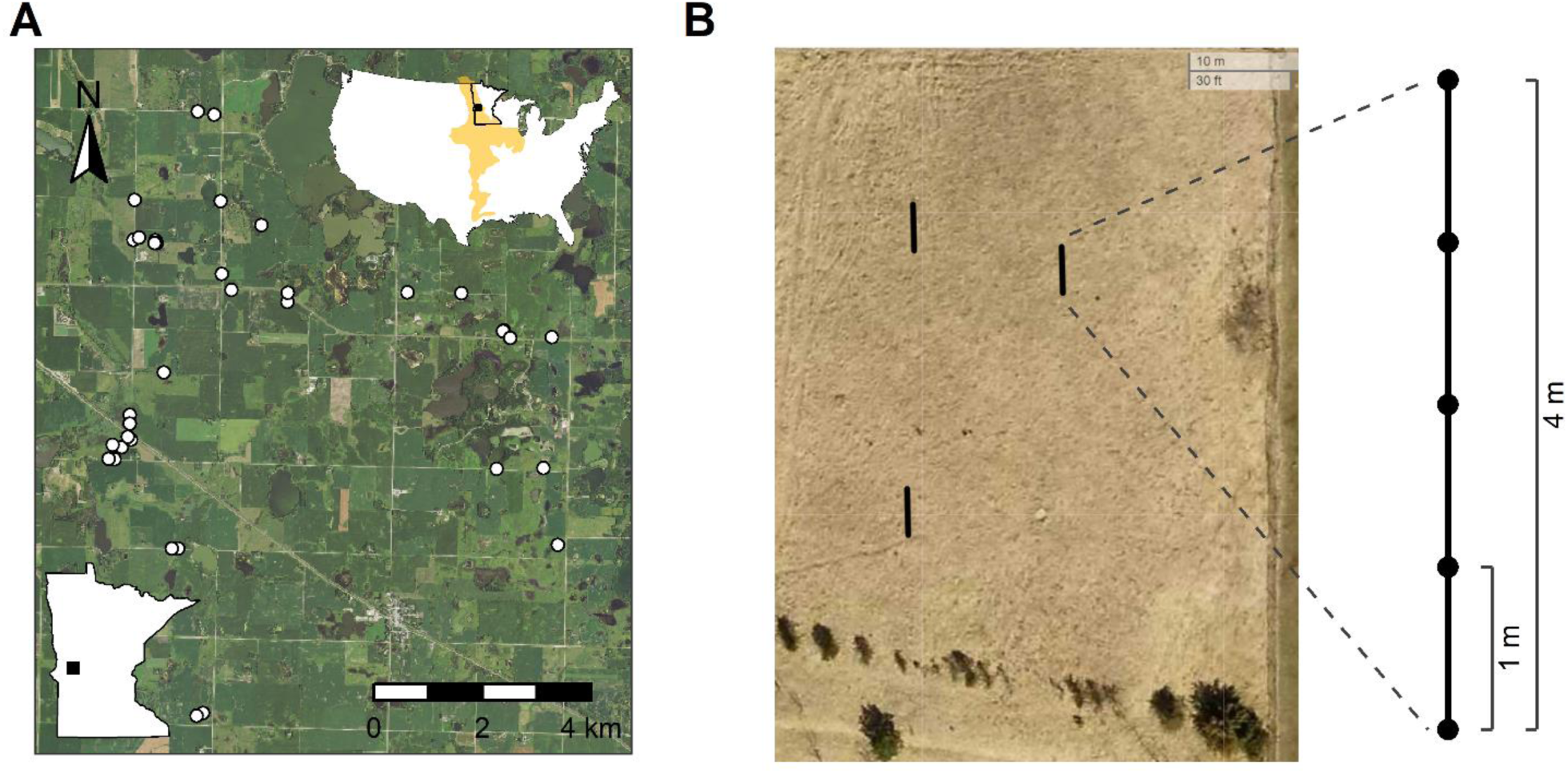
Map depicting the location of study sites in western Minnesota, USA (A) and diagram illustrating placement of experimental transects within a site (B). In (A), white circles illustrate the N = 36 extant *E. angustifolia* populations included in our seed addition experiment. The upper right inset depicts the study location (black square) in relation to the historic extent of tallgrass prairie (shown in yellow). The lower left inset depicts location of study site within the state of Minnesota. In (B), aerial photo imagery illustrates the random placement of experimental transects within one of our study sites. The right inset depicts transect layout. Each transect includes four 1 m segments terminating with nails. We randomly selected one segment to be sown in fall 2022 and one segment to be sown with seeds in fall 2023. The randomly placed transects comprise an unbiased sample of locations that represents conditions experienced during seed dispersal and establishment in natural populations (Table 1). See Table S1 for information of study sites and Fig. S1 for summary of experimental burn treatments.

We conducted 27 prescribed burns over five springs (2020 - 2024) to experimentally generate a gradient of time since fire with five levels: 0, 1, 2, 3, and >=4 years since fire. These treatment levels represent the range of possible biological scenarios for natural seed dispersal and emergence with dormant-season spring fires. For example, the 0-years-since-fire treatment reflects seeds that are dispersed the fall prior to a spring burn (i.e., fire occurs 5-6 months after seed dispersal but 1-2 months prior to seedling emergence). And the 1-year-since-fire treatment reflects seeds dispersed the fall after a spring (i.e., fire occurs 5-6 months before seed dispersal and 12-14 months before seedling emergence). Sowing seed in two fall seasons (2022 and 2003), yielded a design with spatial and temporal replication of burn treatments (Table S1). We note a wildfire occurred in one study site during spring 2020. We treated this wildfire the same as our experimental prescribed fires because it occurred at the same time and in similar conditions to experimental prescribed burns. Two sites were burned in spring 2020 (one the aforementioned wildfire), seven sites in spring 2021, seven sites in spring 2022, six sites in spring 2023, and six sites in spring 2024 (Table S1; Fig. S1).

### Seed addition procedures

In fall 2022, we harvested 155 *E. angustifolia* seed heads from a recently burned site within the study area, “nice” (Table S1). This site was selected because we expected synchronized post-fire reproduction would yield many seed heads and high pollination rates (Wagenius et al. 2020). Moreover, seed harvest here would not interfere with ongoing, long-term demographic research in other sites. We separated achenes from chaff and pooled all ca. 20,000 *E. angustifolia* achenes. We then used a seed blower to exclude light achenes, which are unlikely to contain an embryo, thus obtaining a batch of fruits with high proportion of seeds. We separated fruits into batches of 50, placed them in labeled coin envelopes, then randomly assigned a unique code to each envelope. We X-rayed all envelopes using a low dose that does not affect seed viability (KUBTEC XPERT 80 digital radiography system). Using radiographs, we counted the number of achenes containing embryos, i.e. seeds, in each envelope. In lab trials of randomly sampled envelopes, more than 80 percent of seeds (i.e., achenes containing embryos) germinated.

Coin envelopes were assigned at random to planting locations and sowing years. Most 1 m long segments within experimental transects were assigned a single coin envelope – our low density sowing treatment. However, we assigned two coin envelopes to one randomly selected segment per site in each year – our high density sowing treatment. This allows us to test how the density of sown seeds influences emergence rates and allows us to manipulate variation in seedling density independently of local environmental conditions (Fig. S2). We sowed seeds on 9 November 2022 and 15 November 2023. We sowed a total of 11,435 achenes of which 96.7 percent contained intact embryos (i.e., 11,057 seeds sown).

### Field survey

Each spring (June 6-16 in 2023 and June 8-13 in 2024), we searched experimental transects for newly emerged *E. angustifolia* seedlings. The presence of cotyledons during these early searches allowed us to distinguish newly emerged experimental seedlings from juvenile *E. angustifolia* plants. To track individual seedlings, we placed a colored toothpick 2 cm North of each individual and mapped its location to the nearest 0.5 cm. High-resolution mapping was accomplished by using distance along the 1 m segment as well as perpendicular distance and direction from transect. The combination of colored toothpicks and high-precision spatial maps allowed us to accurately and efficiently track individual seedlings and quantify local conspecific seedling density.

We revisited each transect during August 7-16 in 2023 and August 7-12 in 2024 to score survival from emergence through the first growing season. Individual seedlings were tracked using the unique combination of toothpicks and mapped location. In total, we marked and monitored N = 278 seedlings that emerged in 2023 and N = 696 seedlings that emerged in 2024 (Table 1). All N = 974 seedlings were included in analyses of seedling emergence. Our analyses of seedling survival included only 965 seedlings because four transects were mowed during mid-summer 2024; we retained them in analyses of seedling emergence but excluded them from analyses of seedling survival. One transect was destroyed by heavy machinery during spring 2024. Our analyses of seedling emergence and seedling survival only included 2023 data from this transect.

**Table 1.** Summary of sample sizes for each year and time-since-fire treatment in our seed addition experiment. See Table S1 for detailed site information. All fires occurred in the spring and sowing in November (fall). Note that 0 indicates that fires were conducted the spring after fall sowing (5 months) and 1 indicates fires were conducted one spring before (9 months)

| Years since fire | Sites | Transects | Seeds sown | Seedlings emerged |
| --- | --- | --- | --- | --- |
| 2023 |  |  |  |  |
| 0 | 6 | 11 | 774 | 12 (1.6%) |
| 1 | 5 | 13 | 619 | 60 (9.7%) |
| 2 | 7 | 16 | 1108 | 60 (5.4%) |
| 3 | 2 | 4 | 277 | 21 (7.6%) |
| 4+ | 17 | 40 | 2686 | 125 (4.7%) |
| 2024 |  |  |  |  |
| 0 | 6 | 12 | 716 | 2 (0.3%) |
| 1 | 6 | 10 | 630 | 155 (24.6%) |
| 2 | 4 | 10 | 676 | 60 (8.9%) |
| 3 | 6 | 14 | 917 | 120 (13.1%) |
| 4+ | 16 | 37 | 2559 | 359 (14.0%) |

### Environmental covariates

To assess the potential for biotic and abiotic factors to influence seedling emergence and survival, we quantified six environmental covariates in our transects. In contrast to our experimentally manipulated years since fire treatments, which enable us to infer causal relationships, we can only assess correlations between covariates and seedling fitness. We chose to quantify a suite of environmental covariates that have been hypothesized to influence seedling fitness including light availability, litter depth, soil compaction, slope, aspect, and the local density of conspecific seedlings.

Light availability is a key factor expected to influence seedling performance in grasslands (Leach and Givnish, 1996; Hautier et al., 2009; Alstad et al., 2016;). We measured both litter depth and light availability directly both years (June 6-12, 2023 and June 4-13, 2024). We measured litter depth to the nearest cm at the midpoint of each 1 m segment. The buildup of dead plant material, especially slowly decomposing grass litter, can form a thick layer that intercepts light and may stifle the growth of newly emerged seedlings. We also measured photosynthetically active radiation (PAR) directly using Apogee MX-300 light meters (Apogee Instruments Inc., Logan, Utah, USA). Due to the many sampling locations across many sites, we chose to employ a metric that quantifies the relative amount of PAR reaching ground level by collecting paired PAR measurements: 1) a ground level measure gauging light available to seedlings, and 2 a reading one meter above ground of the light available above canopy at the same time. These paired measurements avoid complications associated with fluctuations in absolute PAR among and within days. We divided observed PAR at 2.5 cm by PAR observed at 1 m above the soil surface. In a pilot study designed to inform our light measurement protocol, we found this proportion was more robust to variation in cloud cover, time of day, and other factors that influence the absolute amount of light reaching ground level. However, the proportional and raw PAR measurements were both sensitive to the time of day and the angle of the sun. Thus, all light measurements were taken within 3 hours of solar noon (10:00 AM - 04:00 PM local time) when PAR measurements were most consistent.

Various soil properties may influence seedling performance. In the study area, one of the most conspicuous edaphic characteristics is the variation from compacted, gravelly soils to rich, uncompacted loams. Compacted soils may impede the growth of seedling roots and prevent seedling establishment (Wernerehl & Givnish, 2015). Within each transect, we measured soil mechanical impedance between 0-10 cm depth using a SpotOn Digital Soil Penetrometer (Innoquest Inc., Woodstock, Illinois, USA) for each of the four 1 m long segments. Measurements were taken at the midpoint of each 1 m segment. All soil compaction measurements were collected over three days (May 24-26, 2023), during which and for the prior one week there was no precipitation.

Physiographic variation such as slope and aspect may be associated with many factors influencing seedling fitness. Steep slopes may contribute to water runoff, leaching, and erosion. In the northern hemisphere, southwest facing aspects receive the greatest sun exposure corresponding to the hottest and driest conditions with relatively high rates of evapotranspiration. Northeast facing aspects receive the least solar radiation. These factors often lead to conspicuous differences in vegetation and environmental conditions. We quantified the slope and aspect at the midpoint of each segment using a 1 m resolution digital elevation model. We calculated slope using an 8-neighbor approach (a 3 x 3 m area). We converted raw estimates of aspect (i.e., compass bearing) to an aspect index (hereafter referred to as aspect) quantifying the absolute value of the difference between compass bearing (in degrees) and southwest aspect (compass bearing of 270 degrees): aspect index = |bearing - 270|. This angular distance metric ranges from 0 when the observed aspect is southwest to 360 when the observed aspect is northeast.

Finally, we calculated the local density of conspecifics for each seedling based on the individuals present during June seedling surveys. Negative density-dependent seedling survival (i.e., declines in per capita seedling survival as the local density of conspecific seedlings increase) has the potential to offset increased seedling emergence. *Echinacea angustifolia* seedlings produce a single leaf and a deep taproot with laterally branching roots. As a result, seedlings can be expected to interact more strongly with nearby neighbors than more distant neighbors. Thus, we calculated a distance-weighted conspecific density metric (*cd_f_*) following Richardson et al. (2024).

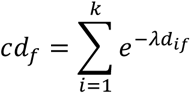

Where *k* is the number of conspecific seedlings within 15 cm of the focal seedling (*f*), *d_if_* is the pairwise distance between seedling *i* and the focal seedling (*f*), and *λ* quantifies the rate at which the influence of seedlings on one another declines with distance. To evaluate potential values of *λ*, we compared the performance of models predicting the probability of seedling mortality as a function of conspecific density using values of *λ* between 0 and 1. Based on this analysis (see Fig. S3), we conducted analyses using distance-weighted measures of conspecific density using *λ = 0*. At *λ =* 0, the *cd_f_* metric simply represents the count of conspecific seedlings within 15 cm of each focal seedling.

### Data analysis

We analyzed seedling emergence using generalized linear mixed effects models (GLMMs). We first modeled the proportion of sown seeds (full achenes) that emerged as a function of years since fire and year sown (2023 or 2024). We treated years since fire as a categorical predictor with five levels: 0 (burned spring after seeds sown), 1 (burned spring before seeds sown), 2 (burned two springs before seeds were sown), 3 (burned three springs before seeds were sown), and 4+ (no burns conducted for at least 4 years). In addition to these fixed covariates, we also included random effects accounting for residual variation among transects and sites (transect nested within site). These random effects account for non-independence among transects within sites that experience the same burn treatment and avoid issues associated with pseudoreplication (Hurlbert, 1984). We modeled seedling emergence using a binomial distribution. We tested the statistical significance of fixed predictors (i.e., year and time since fire) using likelihood ratio tests comparing the full model to reduced models excluding the predictor of interest.

To assess the relationship between seedling emergence and environmental covariates, we also modeled seedling emergence as a function of light availability, soil compaction, slope, aspect, and the density of seeds sown. In this analysis, we excluded data from the treatment 0 years since fire, in which there was consistently low emergence presumably reflecting direct mortality caused by fire (Wagenius et al., 2012). We first examined correlations among these covariates. Two measures of light availability, litter depth and the proportion of PAR reaching ground level, were strongly correlated (Pearson’s correlation: r = -0.53, N = 656, P < 0.001; Fig. S4). To avoid issues associated with collinearity, we chose to include only litter depth in our environmental covariates model. Litter depth measurements were available for every segment and time period (three transects lacked PAR measurements in 2024) and litter depth was more strongly associated with emergence. We applied a log transformation after adding 1 to all litter depth values to improve the distribution of residuals in this and subsequent analyses utilizing litter depth. We fit a full model that included sowing density (high vs. low), year, litter depth, soil compaction, slope, and aspect as fixed predictors and random effects for transect nested within site. We then performed model selection using backwards elimination based on AIC scores (removing terms for which ΔAIC < 2).

To assess fire effects on the seedling survival to late summer (>2 months after emergence), we used GLMM to fit the probability of individual seedling survival to late summer using a Bernoulli distribution. This model included fixed predictors for experimental treatments years since fire and year, as well as random effects for transect nested within site. Due to the paucity of seedlings present in 0 years since fire treatment (N = 14 total seedlings over 2-yr experiment) and lack of variation (all 14 seedlings survived to late summer), we chose to combine the 0 and 1 years since fire treatments for this specific model. Tests were conducted as above to assess the relationship between seedling survival and environmental covariates, including litter depth, soil compaction, slope, aspect, and conspecific density as fixed predictors as well as random effects for transect nested within site. We applied a log-transformation to litter depth and conspecific density after adding 1 to all values. We performed model selection using backwards elimination.

To evaluate how experimental treatments influenced light availability, we analyzed years since fire effects on both the proportion of PAR reaching ground level and litter depth. We fit a GLMM using a beta distribution because the response variable is a proportion constrained between 0 and 1. This model included fixed predictors for experimental treatment and year as well as random effects for transect nested within site. We fit a linear mixed effects model (LMM) to analyze fire effects on litter depth. Our litter depth model also included fixed predictors for experimental treatment and year as well as random effects for transect nested within site. Statistical tests were conducted as above

All analyses were conducted using R (R Core Team, 2024). Statistical models were fit using the glmmTMB package (Brooks et al., 2017) and figures were made using ggplot2 (Wickham, 2016). All data and code necessary to replicate figures and analyses have been archived in a publicly accessible digital repository (Beck and Wagenius, 2024).

## RESULTS

### Seedling emergence

Seedling emergence in our experimentally sown segments varied from 0 to 52 percent (Fig. 2). The experimental years since fire treatments influenced seedling emergence (Table S2; Likelihood ratio test: ΔAIC = 165.9, χ^2^ = 173.86, df = 4, P < 0.001). Seedling emergence also differed between years (ΔAIC = 108.2, χ^2^ = 110.20, df = 1, P < 0.001). Less than one percent of seeds sown prior to burning emerged the following June (estimated percent emerged [95% confidence interval]: 0.4% [0.2 – 0.8] in 2023 and 1.1% [0.6 – 2.1] in 2024), whereas considerably higher proportions of seeds sown in the fall after fire the previous spring emerged (2023: 7.3% (4.8 – 10.8); 2024: 16.5% (11.0 – 24.2)). Seedling emergence was lower 2 years after fire (3.2% [2.1 – 4.8] in 2023 and 7.7% [5.2 – 11.4] in 2024), 3 years after fire (2.6% [1.7 – 4.0] in 2023 and 6.4% [4.3 – 9.3] in 2024), and 4 years after fire (3.9% [2.7 – 5.7] in 2023 and 9.4% [6.8 – 12.9] in 2024). Notably emergence more than four years after fire was highly variable, ranging from 0 to 44% emergence.

**Fig. 2.**
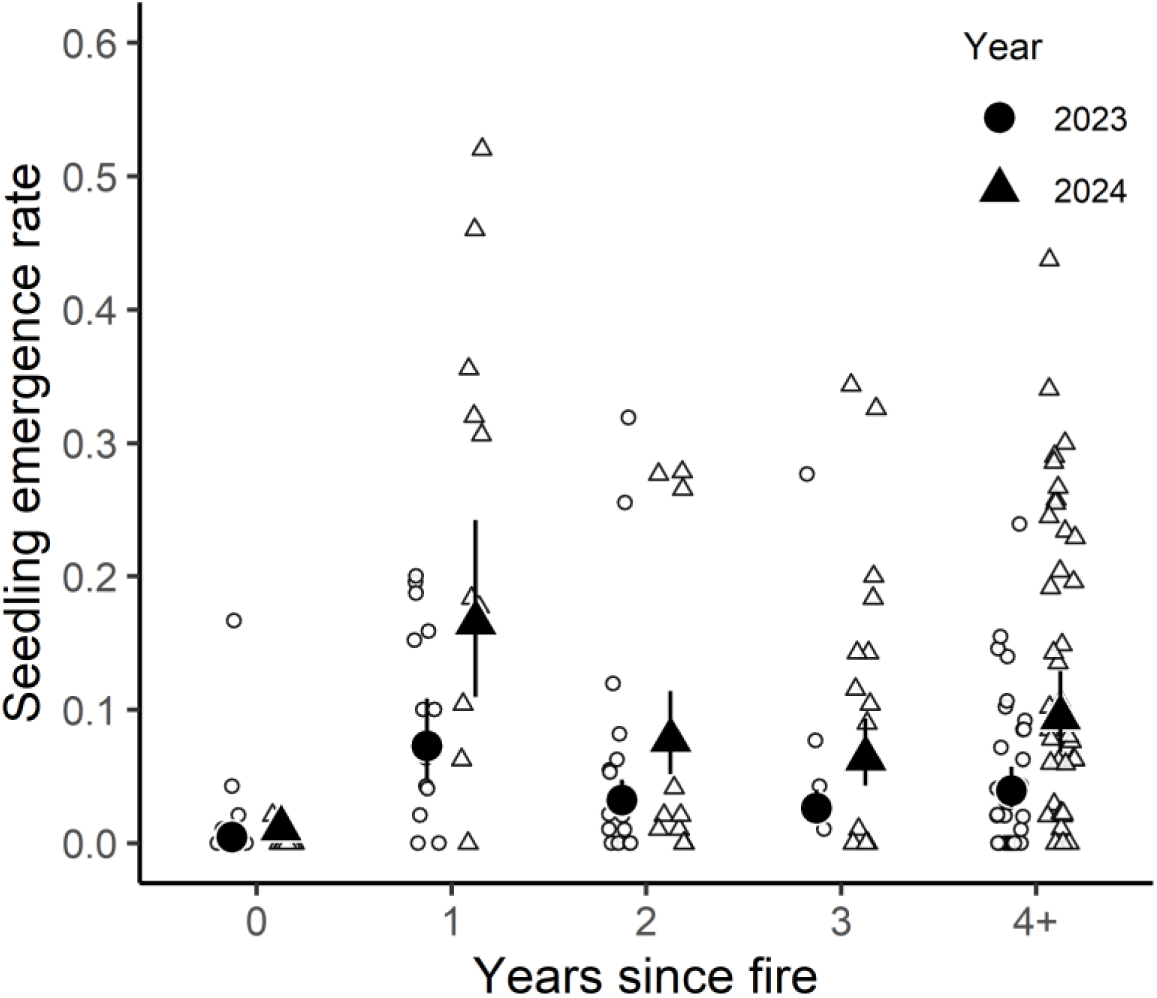
Time since fire influences rates of *E. angustifolia* seedling emergence. Solid black points illustrate predicted mean emergence rates for each experimental treatment and year (see Table S2 for model selection). Small open points illustrate observed emergence per segment within transects (N = 84 transects and N = 167 segments).

In addition to these experimental fire effects, several environmental covariates were associated with variation in seedling emergence (Fig. 3). Across experimental treatments (excluding 0 years since fire), segments sown at high seed densities exhibited lower emergence than low density seeding treatments (ΔAIC = 6.4, χ^2^ = 8.39, df = 1, P = 0.004). Seedling emergence was also negatively associated with litter depth (ΔAIC = 9.4, χ^2^ = 11.38, df = 1, P < 0.001). We found no evidence that soil compaction (ΔAIC = 0.3, χ^2^ =2.28, df = 1, P = 0.131), slope (ΔAIC = 0.4, χ^2^ = 2.36, df = 1, P = 0.124), or aspect (ΔAIC = 1.9, χ^2^ = 0.07, df = 1, P = 0.788) were associated with seedling emergence (Table S3).

**Fig. 3.**
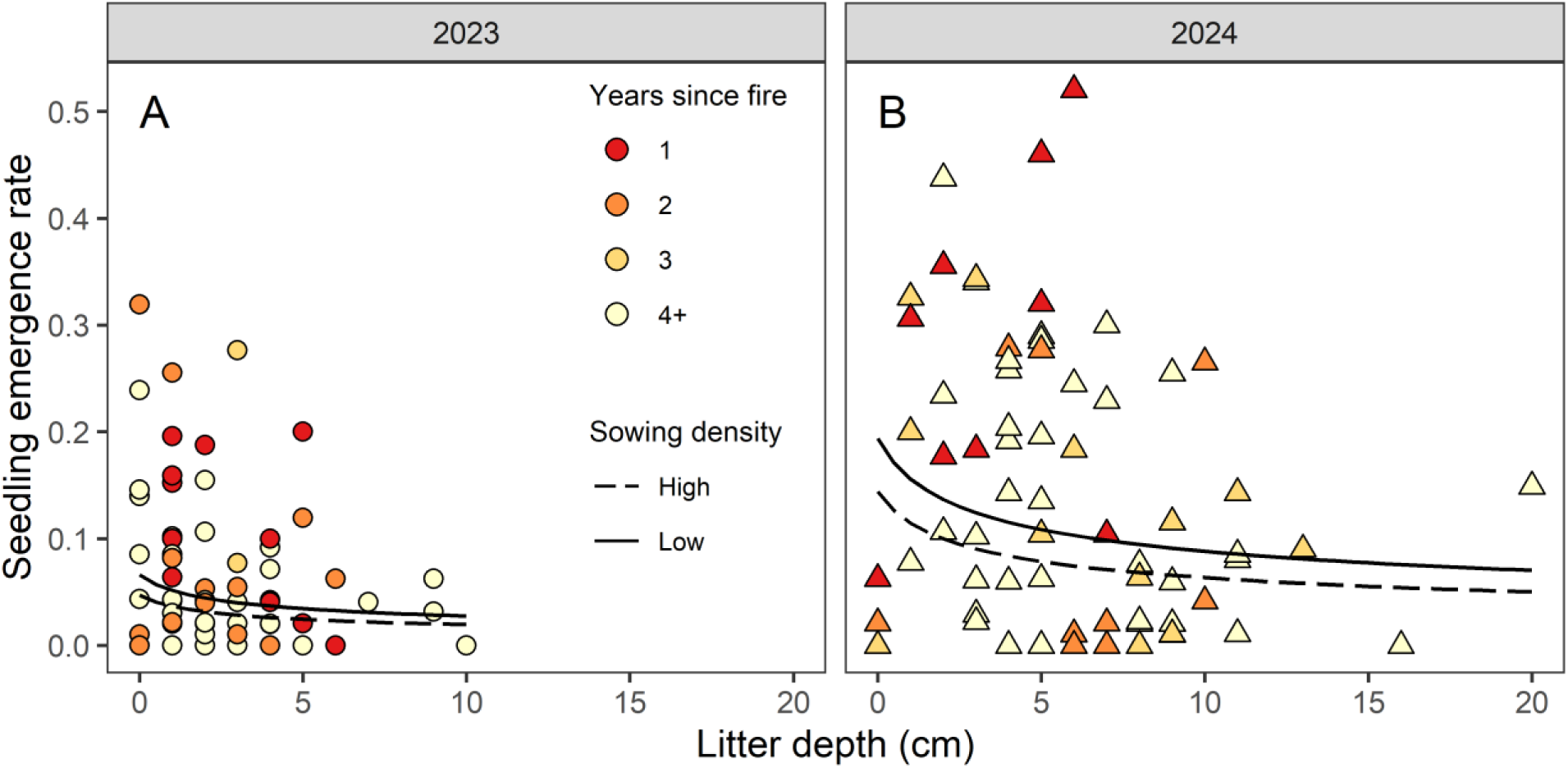
Seedling emergence rates decline with increases in leaf litter, declined in the experimental high-density sowing treatment, and were lower in 2023 (A) compared to 2024 (B). Solid and dashed lines show predicted emergence for the low- and high-density sowing treatment, respectively. Small points depict observed emergence within transects (i.e., number of newly emerged seedlings found divided by number of seeds sown, N= 84 and 83 transects in 2023 and 2024 respectively) and are colored by experimental burn treatment. Note that litter depth, year, and sowing density were retained in the best performing statistical model while slope, aspect, and soil compaction were excluded (see Table S3 for model selection details). Also note that the 0 years since fire treatment was excluded due to direct mortality caused by fire.

### Seedling survival to late summer

Eighty-eight percent (845 of 965) of newly emerged *E. angustifolia* seedlings survived to late summer including 92 percent (257 of 278) in 2023 and 86 percent (588 of 687) of seedlings in 2024 (Fig. 4). Seedling survival varied from 100 percent survival (14 of 14 seedlings) in 0 years treatment to 79 percent (110 of 140) in 3 years since fire treatment. Survival from emergence to late summer differed between years (Table S4; ΔAIC = 5.9, χ2 = 7.874, P = 0.005) and among experimental treatments (ΔAIC = 3.5, χ2 = 9.483, P = 0.024). Survival was highest in the combined 0 and 1 years since fire treatments (predicted probability of survival [95% confidence interval]: 0.97 [0.93 – 0.99] in 2023, 0.93 [0.85 – 0.97] in 2024). Survival declined in the 2 years after fire treatment (0.95 [0.87 – 0.98] in 2023, 0.89 [0.75 – 0.96]) and 3 years since fire treatment (0.86 [0.70 – 0.94] in 2023, 0.72 [0.53 – 0.85] in 2024). Survival was greater in the 4+ years since fire treatment (0.94 [0.87 – 0.97] in 2023, 0.86 [0.78 – 0.92] in 2024), though survival varied greatly among sites and transects within sites in this treatment (Fig. 4).

**Fig. 4.**
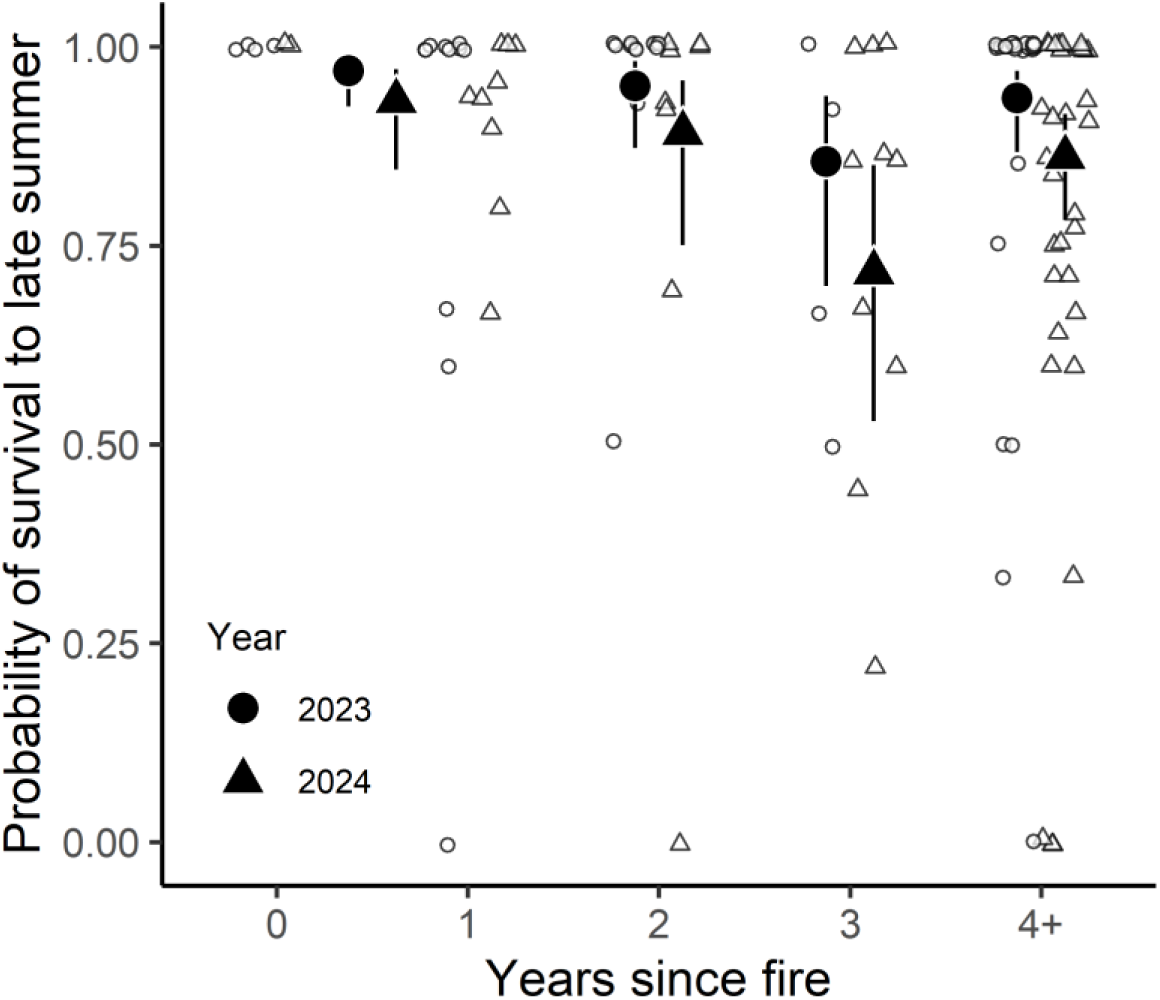
Time since fire influences survival of *E. angustifolia* seedlings. Solid black points illustrate model-predicted survival rates for each treatment and year combination (based on N = 965 *E. angustifolia* seedlings). Small open points illustrate observed survival within transects (N = 84 transects). Note that we combined the 0- and 1-year treatments in our analysis due to the small sample size and lack of variation in the 0 years since fire treatment. See Table S4 for statistical summary.

Several covariates were associated with survival in our analysis encompassing all treatments (Fig. 5; Table S5). The probability of seedling survival was negatively associated with litter depth (ΔAIC = 4.86, χ2 = 6.886, P = 0.009) and positively associated with the local density of conspecific seedlings (ΔAIC = 4.63, χ2 = 6.658, P = 0.010). We found no evidence that seedling survival was associated with soil compaction (ΔAIC = 1.81, χ2 = 0.19, P = 0.662), slope (ΔAIC = 1.98, χ2 = 0.02, P = 0.875), or aspect (ΔAIC = 1.56, χ2 = 0.44, P = 0.509).

**Fig. 5.**
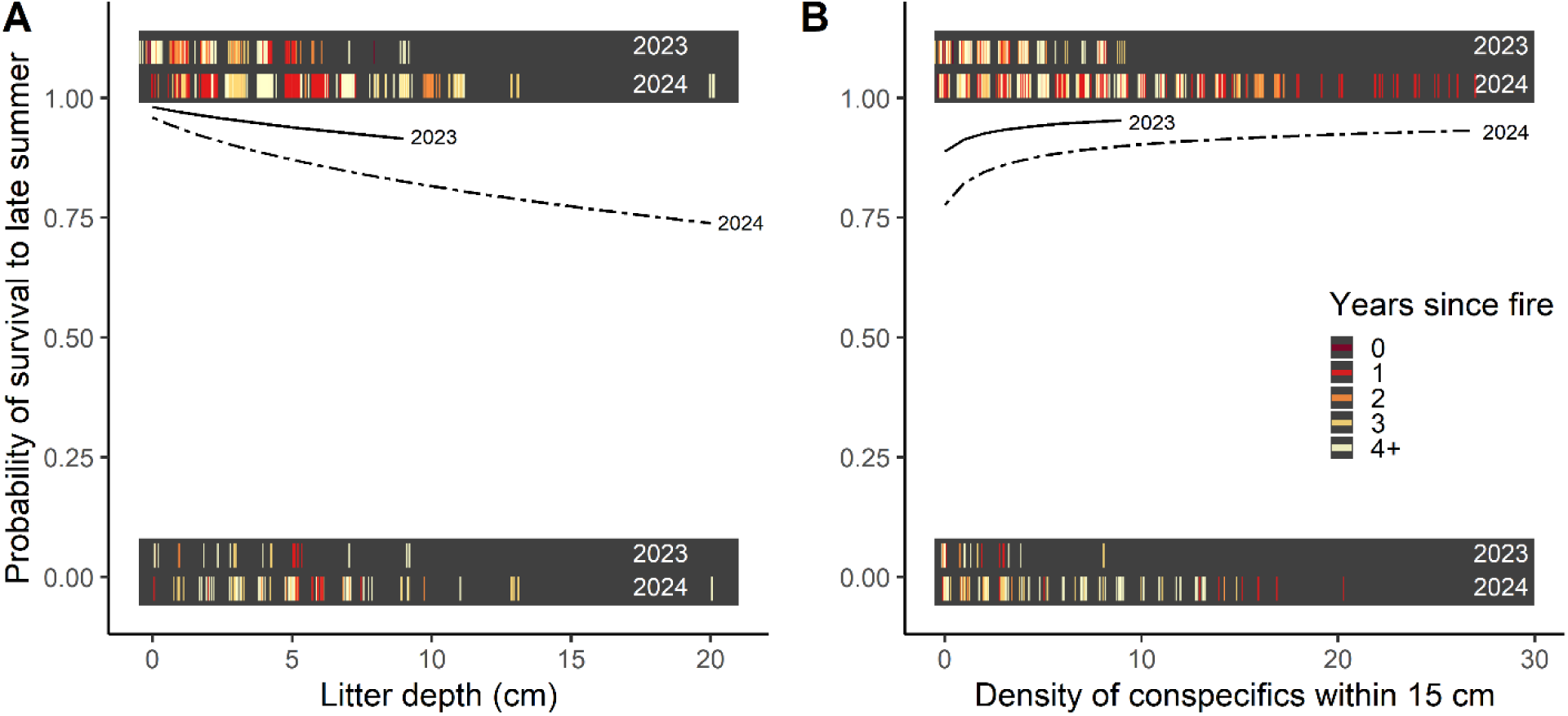
Seedling survival to late summer declines with litter depth (A), increases with density of conspecific seedlings (B), and was greater in summer 2023 than summer 2024. Lines depict predictions from the best supported survival model (Table S3), the solid line depicts predicted survival rates from 2023, and the dashed line depicts predicted survival rates from 2024. Colored tick marks at probability 1 and 0 depict observed survival and mortality events, respectively. The upper and lower rows of ticks in each shaded box corresponds to observed survival and mortality events from summer 2023 and 2024 respectively (N = 965 total *E. angustifolia* seedlings). Boxes were shaded and ticks were jittered to increase visibility. Note that litter depth, year, and conspecific seedling density were retained in the best performing model while slope, aspect, and soil compaction were excluded, see Table S5 for model selection details.

### Fire effects on light availability

Experimental years since fire treatments influenced both measures of light availability (Fig. 6; Table S6). The proportion of PAR reaching ground level depended on years since fire (Fig. 6a; χ2 =140.76, df = 4, P < 0.001) as well as year (χ2 =20.32, df = 1, P < 0.001). PAR reaching ground level was greater in 2023 across experimental treatments and peaked in the 0 years since fire treatment (0.77 [0.71 – 0.82] in 2023 and 0.70 [0.64 – 0.76] in 2024). PAR levels then attenuated and remained fairly consistent across the remaining experimental treatments: 1 year since fire (0.43 [0.35 – 0.51] in 2023 and 0.35 [0.28 – 0.43] in 2024), 2 years since fire (0.40 [0.33 – 0.48] in 2023 and 0.33 [0.26 – 0.40] in 2024), 3 years since fire (0.42 [0.33 – 0.52] in 2023 and 0.35 [0.27 – 0.44] in 2024), and 4+ years since fire (0.45 [0.39 – 0.52] in 2023 and 0.37 [0.31 – 0.44] in 2024). Likewise, litter depth depended on years since fire (Fig. 6b; χ2 = 184.58, df = 4, P < 0.001) and differed between years (χ2 = 50.10, df = 1, P < 0.001). Litter depth was greatest in the 4+ years since fire treatment (3.0 [2.5 – 3.7] cm in 2023 and 4.6 [3.8 – 5.5] cm in 2024) and lowest in the 0 years since fire treatment (0.3 [0.1 – 0.6] cm in 2023 and 0.8 [0.5 – 1.2] cm in 2024). The remaining treatments had intermediate litter depth: 1 year since fire (2.7 [2.0 – 3.4] cm in 2023 and 4.1 [3.2 – 5.1] cm in 2024), 2 years since fire (2.5 [1.9 – 3.1] cm in 2023 and 3.8 [3.0 – 4.7] cm in 2024), and 3 years since fire (2.1 [1.5 – 3.0] cm in 2023 and 3.3 [2.5 – 4.4] cm in 2024).

**Fig. 6.**
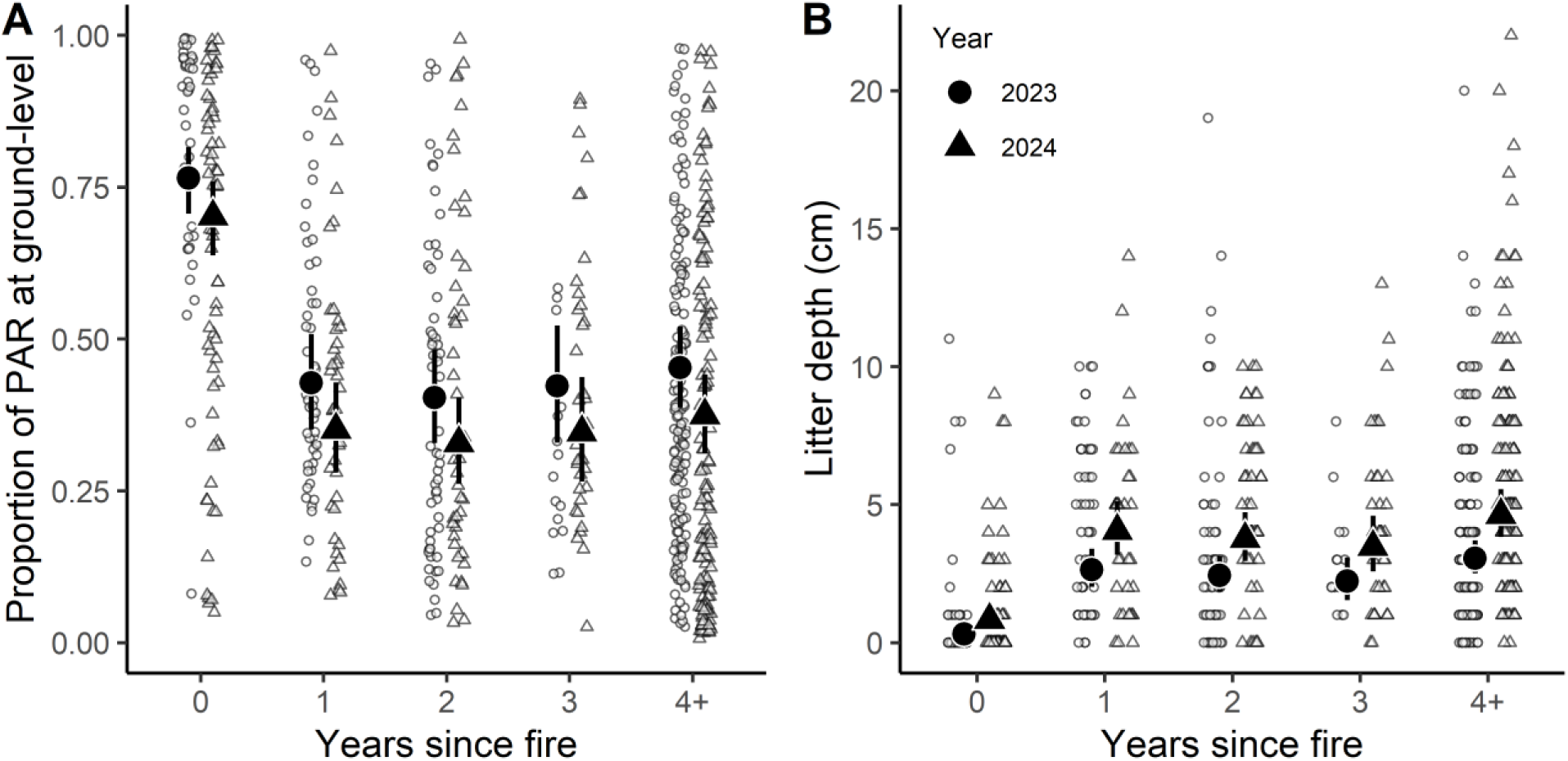
Time since fire influences (A) the proportion of photosynthetically active radiation (PAR) available at ground level and (B) litter depth. Solid black points and lines depict estimated marginal predicted mean values with 95% confidence intervals for all combinations of year and experimental treatment levels (see Table S6 for statistical summary). Small open points represent observed measurements (N = 656 PAR measurements and N = 664 litter depth measurements from 84 transects and 36 study sites). Observed points segregated by year and then jittered horizontally to increase visibility.

## DISCUSSION

By generating conditions favorable for seedling recruitment, periodic fire is expected to promote the persistence of local plant populations and maintain plant diversity in many historically fire-dependent ecosystems. The positive effects of fire on seedling recruitment could plausibly reflect greater seed production, improved seedling emergence, or increased survival of seedlings after fire necessitating experiments capable of discriminating among these processes. In this study, we experimentally manipulated seed inputs to evaluate how years since fire influenced seedling emergence and seedling survival of *E. angustifolia* across 36 patches of remnant tallgrass prairie. Seedling emergence and survival peaked shortly after fire and generally declined with years since fire. This offers empirical support for predictions about fire’s influence on seedling emergence and survival. Moreover, our results are consistent with hypotheses that light plays an important role in mediating the effects of fire on seedling emergence and seedling survival. Although seedling emergence declined when seeds were sown at high densities, overall recruitment increased with seed inputs suggesting both seed availability and microsite conditions may influence seedling recruitment after fire. Our experimental design overcomes limitations associated with earlier observational and experimental studies. It thus enables broader inferences and mechanistic insights that may apply across natural plant populations to clarify demographic processes by which fire influences population dynamics in historically fire-dependent ecosystems.

Consistent with predictions about the role of periodic fire in seedling recruitment within historically fire-dependent ecosystems (Lamont & Downes, 2011; Leach & Givnish, 1996; Menges & Dolan, 1998; Nordstrom et al., 2021; Satterthwaite et al., 2002), our experimental treatments varying years since fire influenced both seedling emergence and seedling survival. Emergence was lowest among seeds sown the fall prior to spring experimental burns. These findings mirror a previous experiment demonstrating low emergence among seeds sown prior to burning presumably due to fire-induced mortality among seeds or newly emergent seedlings that lack energetic reserves to resprout after being damaged (Wagenius et al., 2012). Given that our experimental burns were conducted before seedlings emerged, we suspect direct fire damage to seeds accounts for the low emergence we observed here. The peak in emergence one year since fire treatment corresponds with observations of natural recruitment (Wagenius et al., 2012; Nordstrom et al., 2021). Meanwhile, seedling survival was greatest shortly after fire. Of the very few seedlings that emerged after a fire from seed sown before experimental burns (0 years since fire treatment), all seedlings (14 of 14) survived. Seedlings established from seed sown in the fall after fire also exhibited high survival (199 of 214 survived in the 1 year since fire treatment). Taken together, our findings about germination and early survival parallel findings from observational studies in other species and study systems (e.g., Menges & Dolan, 1998). Conditions favorable for seedling recruitment are often expected to persist for several years after fire or gradually deteriorate leading to continuous declines in seedling recruitment each successive year since fire (Glenn-Lewin et al., 1990). Emergence and survival tended to decline as years since fire increased, but seedling establishment was similar among the 2, 3, and 4+ years since fire treatments, suggesting the conditions favorable for seedling recruitment in prairies and other grasslands may not last as long as commonly expected.

Light availability is widely assumed to mediate the effects of fire on seedling recruitment in prairies and other historically fire-dependent ecosystems (Lamont & Downes, 2011; Leach & Givnish, 1996; Menges & Dolan, 1998; Satterthwaite et al., 2002). Our findings here are largely consistent with this hypothesis. Litter depth was associated with both rates of seedling emergence and the probability of seedling survival, and was correlated with direct measurements of the photosynthetically active radiation available to newly emergent seedlings. In a factorial experiment manipulating seeding density, water availability, competitors, and fire, Zimmermann et al. (2009) similarly found competitors inhibited emergence, and fire effects appeared to be largely mediated by their effects on competition. Nevertheless, we caution that causal relationships between light and seedling fitness cannot be definitely established without experimental manipulation. The most direct evidence for the importance of light for grassland biodiversity comes from experiments in which supplemental light promotes plant survival and by extension species diversity (e.g., Hautier et al., 2009).

The same cautionary note about causal relationships applies to other associations we observed between covariates and seedling emergence and survival. For example, we observed greater seedling emergence, lower rates of early survival, and reduced light availability (lower PAR at ground level and greater litter depth) in 2024 compared to 2023. It is possible these year effects are related to interannual differences in precipitation. Spring and summer 2024 were substantially wetter than 2023 (April – June precipitation of 326 mm in 2024 compared to 154 mm in 2023, PRISM Climate Group 2024). Greater precipitation may have promoted seedling emergence but also stimulated the growth of cool season grasses – especially the exotic *Bromus inermis* – which consequently reduced light availability and contributed to greater seedling mortality. However, without experimental manipulation of precipitation it is impossible to rule out various factors that differed between years (e.g., winter snowfall, spring temperatures and evapotranspiration, etc.). These fundamental issues of correlation and causation are also exemplified by the positive association between local conspecific density and the probability of seedling survival we observed. This positive association contradicts our *a priori* expectation that conspecific density could be negatively associated with seedling survival due to competition for similar resources or greater exposure to pathogens and predators (e.g., Comita et al., 2014). Even though we experimentally manipulated sowing density, realized differences in conspecific seedling density were confounded with environmental factors that promote seedling emergence. Previous studies of natural recruitment in *E. angustifolia* found no evidence of density-dependent mortality (Richardson et al., 2024). Experimental manipulation of seedling density independent of environmental conditions would provide more robust insights into density-dependent seedling fitness though doing so is difficult due to inherent fitness-environment associations.

Our findings here have implications for both the life history and the demography of long-lived plants in historically fire-dependent ecosystems. Many of the long-lived, iteroparous plant species that dominate historically fire-dependent ecosystems conspicuously increase their reproductive effort after fire (Beck et al., 2024; Fidelis & Zirondi, 2021; Lamont & Downes, 2011), including *E. angustifolia* (Beck et al., 2023; Wagenius et al., 2020). Enhanced seedling recruitment after fire is often cited as a potential fitness advantage associated with fire-stimulated flowering (Araújo et al., 2013; L. Zirondi et al., 2021; Lamont & Downes, 2011). This view directly parallels the environmental prediction hypothesis for masting species which posits that concentration of reproduction in years with favorable conditions for seedling recruitment confers a fitness benefit (Beck et al., 2024; D. Kelly & Sork, 2002; Vacchiano et al., 2021). Our results here illustrate the fitness advantages associated with concentrating seed dispersal after fire and suggest enhanced post-fire seedling recruitment is a plausible adaptive explanation for the prevalence of fire-stimulated flowering. However, we note two factors that may diminish potential fitness benefits. First, we found evidence that seeds sown at high densities emerged at lower rates. Thus, the potential fitness benefits associated with concentrating reproduction after fire may saturate with increases in conspecific density. Second, we observed substantial underlying spatial and temporal variation in seedling emergence and survival. The magnitude of this residual variation compared to the effect sizes associated with fire, especially when considered with work demonstrating that fire effects on seedling recruitment vary with environmental context (e.g., Menges & Hawkes, 1998), suggests selection may not be as strong or consistent as commonly assumed (L. Zirondi et al., 2021; Lamont & Downes, 2011). These factors merit consideration in the context of evolutionary explanations for fire-stimulated flowering (Beck et al., 2024).

Variation in recruitment is expected to strongly influence the demography of the long-lived, iteroparous plant species that dominate in many historically fire-dependent ecosystems (Grubb, 1977). These species typically exhibit low survivorship early in life and relatively high survivorship as plants grow and age. Consequently, population dynamics in long-lived plant species are often sensitive to factors that influence seedling recruitment (Bruna, 2003; Grubb, 1977; Menges & Dolan, 1998; Nordstrom et al., 2021). Demographic studies employing population projection models frequently find that population growth rates are sensitive to variation in seedling recruitment rates and that greater post-fire recruitment often promotes population growth (Menges & Dolan, 1998; Nordstrom et al., 2021; Satterthwaite et al., 2002). Our experiment contributes several important insights relevant to fire effects on plant demography. Most generally, our findings suggest both seed availability and microsite favorability may contribute to improved seedling recruitment after fire in historically fire-dependent systems like tallgrass prairie. Further demographic work is needed to quantify the contributions of seed availability and microsite favorability to plant demographic rates and assess the extent to which the positive effects of periodic fire on population growth and persistence are attributable to improved survival (as hypothesized by Leach & Givnish, 1996) or enhanced seed production (as proposed by Wagenius et al, 2020; Beck et al., 2023). Robust demographic inferences about fire effects will depend on incorporating realistic variation in these different vital rates. For example, Nordstrom et al. (2021) found inferences about the effects of fire on population dynamics were sensitive to variation in seedling recruitment. In scenarios with relatively high recruitment, the beneficial effects of fire on seedling recruitment and juvenile survival led to higher rates of population growth after fire. However, in scenarios with low recruitment, seedling recruitment was less influential than other life stages in determining population growth rates. Our experiment, which encompassed representative locations across a heterogeneous landscape, revealed substantial spatial and temporal variation in emergence and survival independent of experimental fire treatments which may have important demographic implications (Quintana-Ascencio, 2023).

The consistently low rates of seedling emergence in sites burned the spring after sowing (i.e., our 0 years since fire treatment) merit special mention in the context of fire frequency and prairie plant diversity. Previous studies investigating the effects of burn frequency have noted that annual burning tends to erode plant diversity over time (Collins & Calabrese, 2012; Johnson & Knapp, 1995). Several previous studies postulate that annual burning increases the dominance of warm season grasses to the point that they competitively exclude herbaceous species (Benson & Hartnett, 2006; Collins & Calabrese, 2012). Our findings suggest an alternative demographic explanation could account for the negative effects of annual spring burning on plant diversity. Annual spring burning may consistently inhibit seedling emergence and greatly reduce seedling recruitment leading to the long-term erosion of plant diversity as sexually mature plants eventually die without replacement. Future demographic work directly investigating this hypothesis would be of great value and contribute mechanistic insights into how fire frequency influences plant population dynamics and plant diversity in tallgrass prairie and other historically fire-dependent ecosystems.

## CONCLUSION

Our experimental investigation of fire effects in a heterogeneous tallgrass prairie landscape revealed that time since fire influences both seedling emergence and seedling survival. We found evidence supporting the hypothesis that the microsite characteristics, specifically light availability and litter density, mediate fire effects on seedling growth and survival. Further, overall recruitment increased with seed supply. Thus, our experiment reveals that both seed availability and microsite conditions can influence seedling recruitment after fire in natural populations.

## ACKNOWLEDGMENTS

We thank R. Shaw for advice and helpful discussions during the design of the experiment. We thank members of the Echinacea Project – especially M. Stevens, A. Carroll, L. Paulson, W. Mosiman, and A. Widdell – for assistance in the field and in the lab. R. Shaw, A. Iler, and P. CaraDonna generously provided feedback that improved the manuscript. This research was supported by awards from the National Science Foundation (2032282, 2050455, 2051562, 2115309) and the Minnesota Environment and Natural Resources Trust Fund as recommended by the Legislative-Citizen Commission on Minnesota Resources (award 2022-091).

