## Supplemental Materials for "Seedling recruitment after fire: Disentangling the roles of microsite conditions and seed availability"

### 14 SUPPLEMENTAL MATERIALS

Table S1. Information about study sites. For each study site, we summarize burn history since 2018, the number of experimental transects, the number of seeds sown in each year, and the number of *E. angustifolia* seedlings that emerged in each year.

| Site abbrev. | Recent burn history | Transects | Sown (2022) | Emerged (2023) | Sown (2023) | Emerged (2024) |
|---|---|---|---|---|---|---|
| aa | 2018 | 3 | 193 | 6 | 200 | 29 |
| alf | 2023, 2024 | 4 | 236 | 2 | 243 | 12 |
| beng | 2020 | 2 | 133 | 7 | 150 | 33 |
| dog | 2021, 2023 | 2 | 149 | 0 | 144 | 22 |
| eelr | — | 3 | 196 | 10 | 193 | 33 |
| eri | 2021 | 2 | 144 | 15 | 138 | 36 |
| eth | — | 2 | 146 | 2 | 143 | 10 |
| fern | — | 2 | 144 | 14 | 146 | 47 |
| gc | — | 2 | 142 | 2 | 142 | 22 |
| hulze | 2024 | 2 | 146 | 2 | 97 | 1 |
| hulzw | — | 2 | 143 | 1 | 147 | 26 |
| hutche | — | 2 | 142 | 9 | 149 | 6 |
| hutchw | 2024 | 2 | 143 | 10 | 136 | 0 |
| kj | 2021, 2023 | 2 | 144 | 9 | 146 | 15 |
| krus | — | 3 | 192 | 7 | 189 | 8 |
| lce | 2021 | 3 | 194 | 26 | 198 | 53 |
| lcw | 2022 | 3 | 139 | 16 | 192 | 41 |
| lfe | 2021 | 3 | 191 | 8 | 197 | 21 |
| lfw | 2022 | 3 | 141 | 15 | 192 | 16 |
| mapp | 2022 | 2 | 95 | 3 | 146 | 2 |
| nice | 2022, 2024 | 3 | 147 | 21 | 196 | 0 |
| nrrx | — | 2 | 146 | 1 | 148 | 27 |
| nwlf | 2022 | 2 | 97 | 5 | 146 | 1 |
| onts | — | 3 | 190 | 10 | 185 | 17 |
| ri | — | 3 | 187 | 31 | 191 | 38 |
| rrx | — | 2 | 146 | 7 | 143 | 31 |
| sape | 2021 | 2 | 147 | 2 | 143 | 1 |
| sapw | 2021 | 2 | 142 | 2 | 142 | 0 |
| sgc | 2023 | 2 | 148 | 0 | 95 | 39 |
| th | — | 2 | 145 | 9 | 135 | 8 |
| torgen | 2024 | 2 | 144 | 1 | 96 | 1 |
| torges | 2023 | 2 | 143 | 0 | 148 | 67 |
| tower | — | 2 | 144 | 15 | 152 | 21 |
| waa | 2022, 2023 | 2 | 95 | 3 | 48 | 3 |
| yohe | 2021 | 2 | 142 | 4 | 99 | 9 |
| yohw | 2021, 2024 | 2 | 148 | 3 | 143 | 0 |

Table S2. Summary of statistical models quantifying effects of time since fire on seedling emergence. For each term, we present the ΔAIC (difference in AIC between full model and model excluding term of interest) as well as the test statistic (χ2), degrees of freedom (df), and P-values for likelihood ratio tests comparing full model to reduced model excluding the term of interest. All models include random effects for transects nested within sites.

| Model term | ΔAIC | χ2 | df | P-value |
|---|---|---|---|---|
| Years since fire | 165.9 | 173.86 | 4 | <0.001 |
| Year | 108.2 | 110.20 | 1 | <0.001 |

Table S3. Model selection quantifying the association between seedling emergence and environmental characteristics (N = 143 segments, excluding 0 years since fire). We performed stepwise backwards elimination. For each candidate model, we present ΔAIC (difference in AIC between models) as well as χ2 and P-values from likelihood ratio tests. Covariates are presented in the order they were excluded during model selection, in other words, the term excluded in the first step of backwards elimination is shown first. All models included random effects for transects nested within sites. Model terms retained in the best performing model are shown in bold.

| Model term | ΔAIC | χ2 | df | P-value |
|---|---|---|---|---|
| aspect | 1.9 | 0.07 | 1 | 0.788 |
| slope | 0.4 | 2.36 | 1 | 0.124 |
| soil compaction | 0.3 | 2.28 | 1 | 0.131 |
| **Litter depth** | **9.4** | **11.38** | **1** | **<0.001** |
| **Sow density** | **6.4** | **8.39** | **1** | **0.004** |
| **Year** | **103.6** | **105.63** | **1** | **<0.001** |

Table S4. Summary of models quantifying the effects of time since fire on seedling survival. For each candidate model, we present AIC and ΔAIC (difference in AIC between focal model and best fitting model) values. We also include χ2 and P-values for a likelihood ratio test comparing the candidate model to the maximal model. All models include random effects for transects nested within sites. The best fitting model includes both terms.

| Model term | ΔAIC | χ2 | df | P-value |
|---|---|---|---|---|
| Years since fire | 3.5 | 9.48 | 3 | 0.024 |
| Year | 5.9 | 7.87 | 1 | 0.005 |

Table S5. Summary of model selection for statistical models quantifying the association between seedling survival and six environmental covariates. We performed stepwise backwards elimination. For each candidate model, we present ΔAIC (difference in AIC between models) as well as χ2 and P-values from likelihood ratio tests. Covariates are presented in the order they were excluded during model selection, in other words, the term excluded in the first step of backwards elimination is shown first. All models included random effects for transects nested within sites. Model terms retained in the best performing model are shown in bold.

| Model term | ΔAIC | χ2 | df | P-value |
|---|---|---|---|---|
| Slope | 1.98 | 0.02 | 1 | 0.875 |
| soil compaction | 1.81 | 0.19 | 1 | 0.662 |
| Aspect | 1.56 | 0.44 | 1 | 0.509 |
| **litter depth** | **4.86** | **6.886** | **1** | **0.009** |
| **conspecific density** | **4.63** | **6.658** | **1** | **0.010** |
| **Year** | **4.08** | **6.110** | **1** | **0.013** |

Table S6. Summary of statistical models assessing the effects of time since fire on light availability, quantified as the proportion of photosynthetically active radiation – PAR – at ground level, and litter depth.

| Model | ΔAIC | χ2 | df | P-value |
|---|---|---|---|---|
| *Proportion PAR* | | | | |
| Years since fire | 18.4 | 20.40 | 4 | <0.001 |
| Year | 132.9 | 140.85 | 1 | <0.001 |
| *Litter depth (cm)* | | | | |
| Years since fire | 49.1 | 51.1 | 4 | <0.001 |
| Year | 175.8 | 183.84 | 1 | <0.001 |

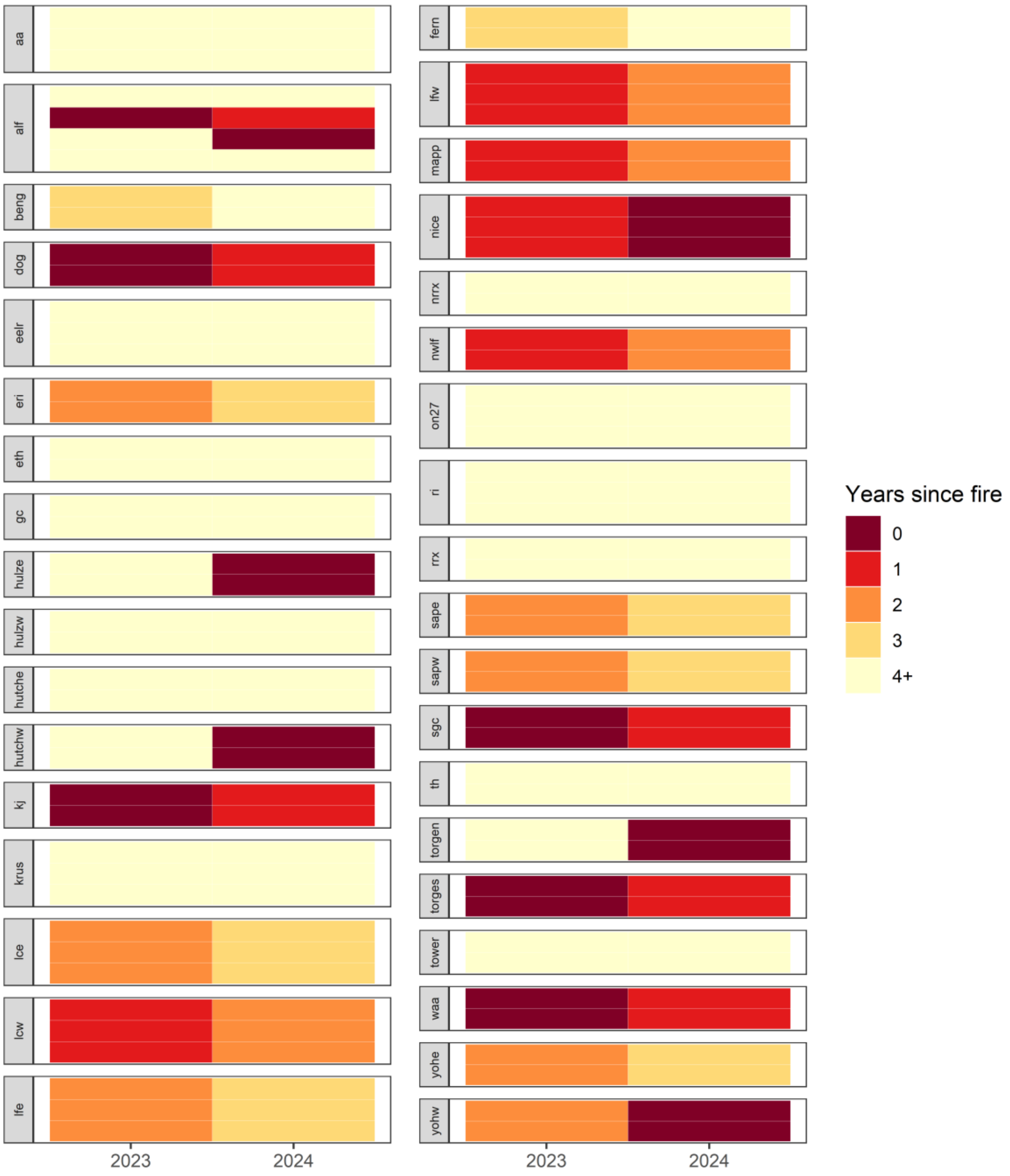


Fig. S1. Heatmap illustrating experimental time since fire gradient across sites and years. Each cell corresponds to a transect-year combination. Cells are grouped by site. Note that there are between 2 and 4 transects per site. The number of transects per site is roughly proportional to site area. Cell colors correspond to experimental time since fire treatment.

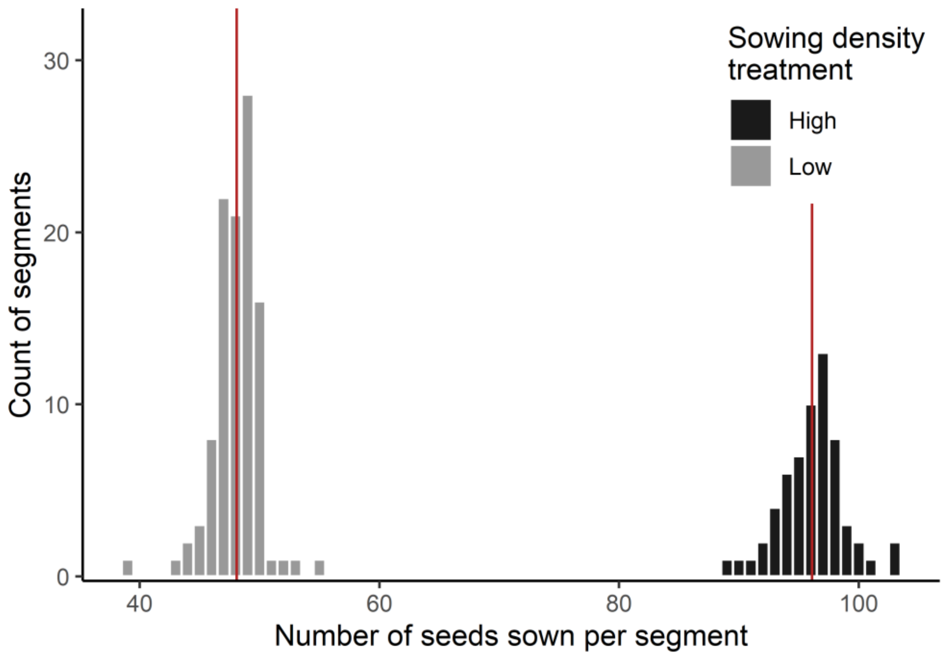


Fig. S2. Histograms of the number of seeds sown per 1 m trasect in the high- and low-density sowing treatments. We varied the density of seeds sown to evaluate how seed availability influenced emergence and to attempt to generate variation in conspecific seedling density independent of environmental conditions. In the lab, approximately 50 fruits were placed in each envelope. The number of seeds per envelope reflects the realized number of achenes with intact embryos sown. See Materials and Methods for details about how seed counts were obtained using X-ray radiographs. The high-density treatment, 2 envelopes, averaged 96.1 seeds per segment (range 89 – 103 seeds) while the low-density treatment, one envelope, averaged 48.1 seeds per segment (range 39 – 55 seeds).

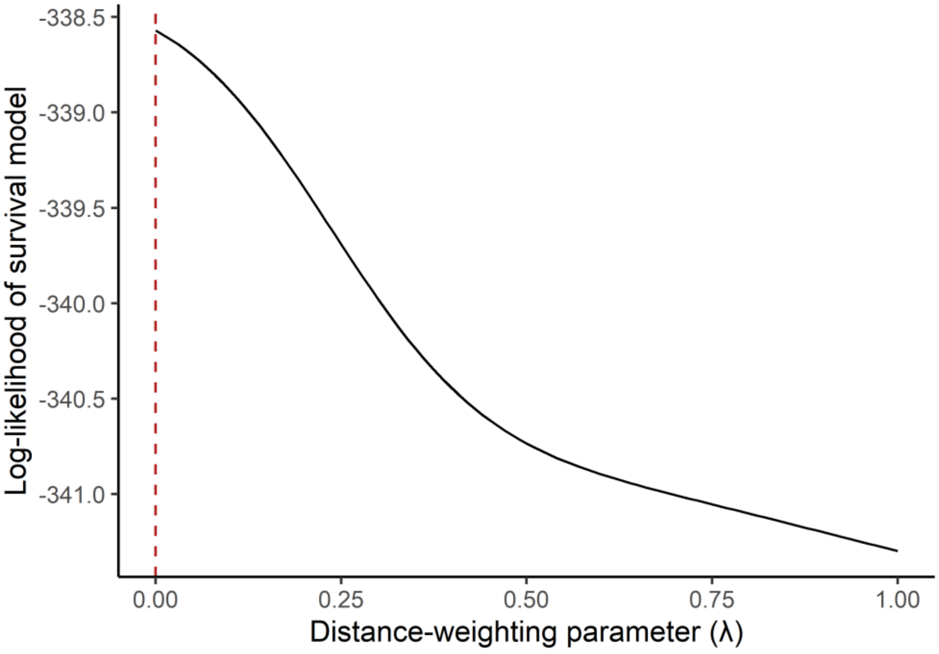


Fig. S3. Profiled log-likelihood of distance-weighting parameter (λ) for statistical models quantifying association between local conspecific seedling density and the probability of seedling mortality. Profiled log-likelihood of λ – rate of exponential decay. The log-likelihood was maximized at λ = 0 (depicted by dashed red line). A value of λ = 0 weights all conspecific seedlings within 15 cm of focal seedling equally.

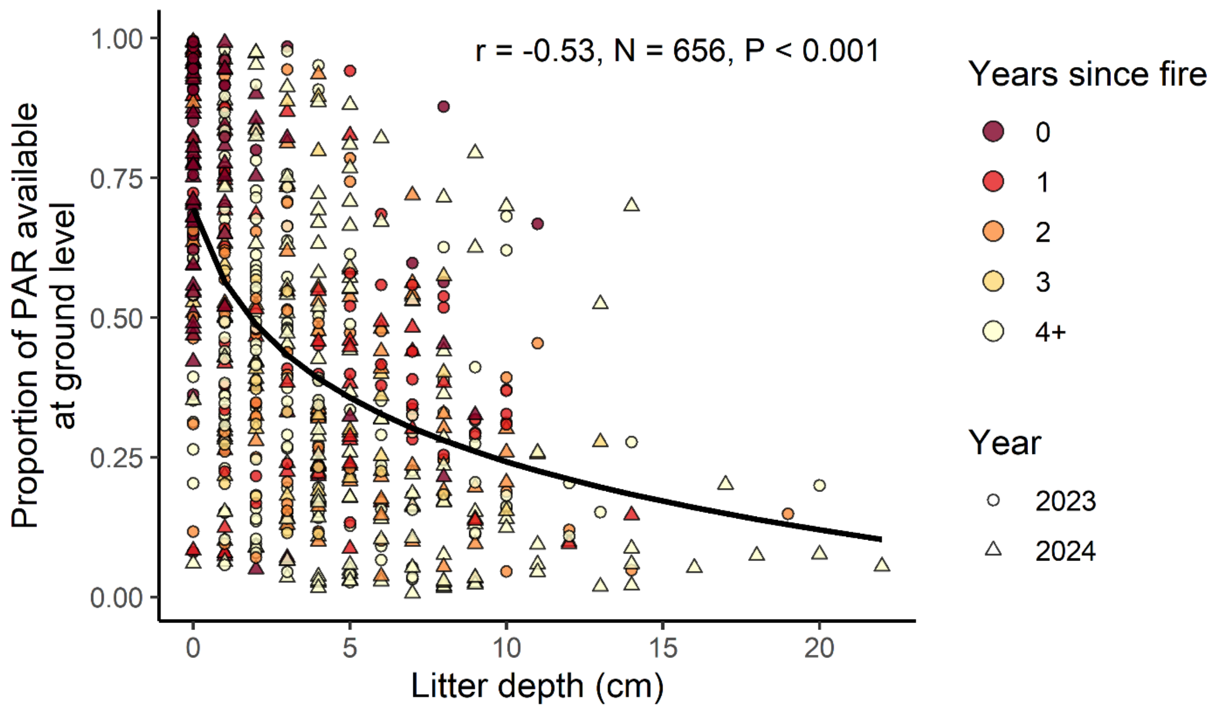


Fig. S4. Litter depth is correlated with the proportion of ambient light reaching ground level (r = -0.53, N = 656, P < 0.001). Note that for statistical analysis we applied a log transformation to litter depth after adding 1 to all values, though the relationship is shown on untransformed x-axis.
